# Brain anatomy in major hormonal transition phases: Longitudinal and cross-sectional volume associations with menarche and menopause

**DOI:** 10.64898/2026.03.31.715492

**Authors:** Melanie Freund, Gloria Matte Bon, Alkistis Skalkidou, Birgit Derntl, Tobias Kaufmann

**Affiliations:** Department of Psychiatry and Psychotherapy, Tübingen Center for Mental Health, University of Tübingen, Tübingen, Germany; Department of Women’s and Children’s Health – Obstetric & Reproductive Health Research, Uppsala University, Sweden; German Center for Mental Health (DZPG), partner site Tübingen, Tübingen, Germany; Centre for Precision Psychiatry, Division of Mental Health and Addiction, Institute of Clinical Medicine, University of Oslo, Oslo, Norway

**Keywords:** female reproductive lifespan, neuroplasticity, menarche, menopause

## Abstract

**Background:** Hormonal transition phases represent windows of increased neuroplasticity across the female lifespan. In this study, we aim to investigate the brain anatomical architecture of hormonal transition phases by directly comparing menarche, as a period of rising levels of steroid hormones, and menopause, as a time of declining levels.

**Methods:** We fit linear models on cross-sectional and linear mixed-effect models on longitudinal magnetic resonance imaging (MRI) datasets, to explore the effects of menarche onset (ABCD study data, N_cross-sectional_=1274, N_longitudinal_=611) and transition into menopause (UK Biobank data, N_cross-sectional_=1614, N_longitudinal_=212) on 66 cortical and 135 subcortical brain volumes, and to identify brain structures with opposing but regional overlapping effects in both periods. Models were adjusted for age and corrected for multiple comparison (P <.05; FDR-corrected).

**Results:** Cross-sectionally, using a between-subject design, 83 brain volumes showed effects of menarche-onset and 17 volumes showed effects of menopause-transition. Of these, seven brain volumes were significantly affected by both transitional periods, showing opposing directional volume changes. Longitudinally, using a within-subject design, 56 brain volumes exhibited menarche effects, of which 46 replicated cross-sectionally. No menopause effect survived correction for multiple comparison, likely due to limited longitudinal sample size.

**Conclusion:** Our findings confirm regionally overlapping brain structural alteration between the two hormonal phases – menarche and menopause – showing the hypothesized opposite effect directions. Additionally, our results show the robustness of menarche effects, which converged across cross-sectional and longitudinal study designs. Taken together, our results contribute to a better understanding of hormone related neuroplasticity, emphasizing the importance of not only understanding individual phases, but understanding the overarching patterns across the female reproductive lifespan.

## Introduction

Across a female’s reproductive lifespan, steroid hormone levels undergo considerable fluctuations. The concentration of oestradiol and progesterone rise with the onset of puberty, fluctuate at generally high levels during the reproductive years across the menstrual cycle and possible pregnancies, and eventually decrease with the transition into menopause ^1^.

Between the ages of eight to fourteen years ^2^ most females enter puberty, the transition from childhood to adulthood, driven by neuroendocrine mechanisms that transform the psychophysiology for maturation and reproduction ^3^. Later in life, over a period of 10 to 15 years, the transition into menopause begins, and around the age of 51 years ^4^ many females have experienced the cessation of menstruation, induced by age-related hormonal processes, leading to the loss of the reproductive capacities ^5^.

These major periods of substantial hormone dynamics experienced by all females with intact ovarian functions, often referred to as hormonal transitional phases, have previously been linked to alterations in brain structure ^6^. Peper et al. found higher levels of oestradiol tied to smaller absolute gray matter volume in females but not in males during puberty ^7^, while Kim et al. found a positive association with oestradiol levels and superior temporal gyrus volume in post-menopausal females ^5^. Pregnancy, another key transitional phase characterized by profound hormonal shifts and experienced by many females, has similarly been linked to cerebral morphometric changes ^8–11^. It has been suggested that changes observed from pre to post pregnancy may resemble changes to those observed from mid-puberty through the following two years in female adolescents, with both marked by gray matter volume reduction alongside rising steroid hormone levels ^12^.

Despite differences in timing, the different hormonal transition phases show notable similarities such as the co-occurrence of rapid hormonal fluctuations and an increased risk of mental disorders ^13^. Higher incidence rates of anxiety and depression in females compared to males are observed during puberty ^14^, and the development of depression and anxiety are linked to peri- and postpartum periods ^15,16^ and menopause ^17^. Directly comparing brain structural changes associated with different transition periods could complement a lifespan picture of common patterns of female neuroplasticity in periods of profound vulnerability.

In the present study we aim to characterize neuroanatomical patterns linked to the transition from before and after menarche and the transition into menopause. We then aim to compare these within-transition pattern to identify shared neural signatures as well as unique, phase-specific patterns of gray matter volume changes.

We used magnetic resonance imaging (MRI) data from the Adolescent Brain Cognitive Development study (ABCD) to investigate effects of puberty, and MRI data of females who entered menopause sourced from the UK Biobank to investigate effects of the transition into menopause, through both, cross-sectional and longitudinal analysis. Our hypothesis was that we would observe similarities between puberty and menopause in the anatomical maps (some of the same regions affected) yet with opposing effect directions given that puberty is commonly associated with a rise and menopause with a decline in sex hormone levels.

## Methods

### Sample Description and Participant Selection

#### ABCD

To investigate female brain changes during puberty, structural MRI data from the Adolescent Brain Cognitive Development (ABCD; release 5.1) study was analysed. This ongoing longitudinal study follows children from approx. 9 years of age into adulthood. Neuroimaging assessments are conducted at 21 scan sites across the United States. The data release contains phenotypic and imaging data of three time points – baseline, 2-year follow-up and 4-year follow-up. In order to assess the pubertal status, participants and their guardians answered questions of the 5 item pubertal development score (PDS) ^18^. All participants and their legal guardians provided written informed consent.

For the analysis, participants were selected in line with the following criteria:

1) Biological sex ‘female’ consistently indicated in both, parents’ and children’s reports.
2) Answer to question “Have you begun to menstruate (started to have your period)?” was not “Decline to answer” or “I don’t know” and matched between parents’ and children’s report.

The longitudinal analysis incorporated MRI data obtained from two consecutive sessions approximately two years apart. The selection of sessions was based on the onset of first menstrual bleeding, also referred to as menarche, specifically including MRI data from the session participants first reported menstruation (PDS question five for females “pds_f5_y” is 4, meaning “yes”, for the first time) and its immediately preceding session (with answer to “pds_f5_y” being “no”). This way, we were able to capture the change in menstrual status from before to after first menstruation in a longitudinal manner. Consequently, we excluded participants reporting menstruation at baseline (as the change in menstrual status happened before the study started) and participants never reporting menstruation over the three sessions – for a total of 611 participants (pre-menarche mean age ± sd = 10.73 ± 0.91 years, post-menarche mean age ± sd = 12.80 ± 0.93 years) (Table 1).

**Table 1.** Demographics overview of the different cross-sectional and longitudinal samples. Pubertal and menopausal ages are given in years (y).

| Analysis design | Menarche<br>ABCD | Menopause<br>UK-Biobank |
| --- | --- | --- |
| cross-sectional | N = 1274 (637 per group) | N = 1614 (807 per group) |
| Mean age | Menarcheal = 12.23 y<br>Non-menarcheal = 12.22 y | Menopausal = 52.93 y mo<br>Non-menopausal = 53.12 y |
| longitudinal | N = 611 | N = 106 |
| Mean age | Pre = 10.73 y<br>Post = 12.80 y | Pre = 51.24 y<br>Post = 55.02 y |

To complement the analysis of within-subject longitudinal change, we also performed a cross-sectional between-subject comparison of two groups of age matched individuals, one pre and one post first menarche. A matching procedure (explained in more detail in section *Matching Procedure*) was applied to the ABCD study data to retrieve two groups with similar age and image quality characteristics, yielding a total of 1274 participants, of which 637 participants reporting menstruation (mean age = 12.23 years, sd = 0.78 years) and 637 not reporting menstruation (mean age = 12.22 years, sd = 0.78 years; Table 1).

Participants from the longitudinal analysis were included into the cross-sectional analysis but only with one session, so they were matched with different individuals rather than their own repeated measurements. The decision not to exclude the participants from the cross-sectional analysis was done to prevent atypical pairings within the matched sample. Participants excluded from the longitudinal analysis mainly represented statistical outliers, either starting their menstruation very early or not initiated menstruation in an older age. Matching these outliers with participants exhibiting age-appropriate development, could compromise the representativeness of the sample and potentially bias the findings with respect to population norms.

#### UK Biobank

To investigate the effects of menopause on the female brain, phenotypic and structural MRI data was accessed from the UK Biobank which is a biomedical database containing biological, lifestyle and health information of half a million participants with ongoing brain MRI data collection at four sites across the United Kingdom. At baseline, the cohort included adults spanning midlife to older adulthood and a subset of individuals has been followed up with a second brain scan.

Our general including criteria consisted of:

1) biological sex is female
2) no hysterectomy or (bilateral) oophorectomy
3) existing imaging data
4) information on menopausal status and age at last period

In line with the procedures for puberty, we also performed two analysis streams for menopause: longitudinal within-subject and cross-sectional group comparison. For the longitudinal analysis, only females attending two consecutive MRI scanning sessions, reporting no menopause in their first session, and reporting menopause in their second session were included. Additionally, we verified the consistency of self-reported age at last period. Participants whose reported age at last menstruation was younger than their age at the first MRI session were excluded from the analysis. This resulted in 106 participants (Table 1) in the pre-menopausal phase (mean age = 51.2 years, sd = 2.6) for which we have follow-up data in the menopausal phase (mean age = 55.0 years, sd = 2.2). Due to the limited sample size, we opted not to impose additional age range criteria, which resulted in an age span of 47 to 55 years at the first session and 52 to 64 years at the second session with a mean time gap between the sessions of 3.78 years (sd = 1.81 years).

For the cross-sectional analysis, participants from the longitudinal analysis were excluded. This decision was based on the considerably larger sample size available for the cross-sectional analysis compared to the longitudinal dataset, which meant that including longitudinal participants would not have added much additional value. In contrast, excluding these participants allowed us to maintain an independent cross-sectional sample.

For most participants, imaging data were available from only a single scan session. In instances where multiple scans existed for a participant, but were not suitable for the longitudinal analysis, due to an unchanged menopausal status across sessions only the data from the initial session remained in the analysis. To avoid potential biases introduced by outliers with very early or late menopause, and to ensure a more representative cross-sectional sample, we restricted the age range to 45 to 60 years participant to be consistent with the literature ^19^.

The matching procedure is explained in detail in section 2.2 and yielded a sample of 1614 participants (Table 1), 807 in the non-menopausal group (mean age = 52.93 years, sd = 2.19 years) and 807 in the menopausal group (mean age = 53.12 years, sd = 2.23 years).

To rule out potential confounds due to menopause hormone therapy (MHT), we performed a supplementary analysis on the menopause dataset, excluding participants who reported MHT use (Supplement Figure 2).

#### Matching Procedure

For the cross-sectional analysis, participants were divided into groups based on their menarche or menopause status, independently of the session.

The groups were then matched on age and image quality (using Euler number as a proxy ^20^ in the menarche data and the head motion variable ^21^ for menopause data).

Participants were divided according to their menarche or menopause status. Each participant in the menarche group was matched with a participant in the non-menarche group that had a Euler number of up to one standard deviation of the left-right average and a difference in age of maximum 2 months (adapted from Wiersch et al., 2023) ^22^. Once matched with one session, the other session of that participant was excluded from the further matching procedure, to ensure participants were only matched once and did not confound group independence.

Similarly, each participant in the menopause group was matched with a participant in the non-menopausal group that had a Euler number of maximal one standard deviation and a difference in age of maximum 12 months.

To achieve the smallest feasible maximum age difference, participants ages were expressed in months, which were directly available from the ABCD study and calculated from participant’s birthdates for UK Biobank. An iterative approach was employed, progressively decreasing the maximum age window in 2-month steps starting at one year for menarche and 2 years for menopause. This yielded a maximum difference of 2 months for menarche and 12 months for menopause, with no significant between-group age difference (t-test, p > 0.05).

We adopted a less stringent age difference among menopause participants compared to menarche participants, as a reflection for the differential rates of brain change across these life stages. Youths’ neurodevelopment is characterized by rapid and substantial structural changes, whereas brain maturation and plasticity slow considerably by mid-adulthood. Therefore, stricter age matching is warranted for participants undergoing menarche to accurately capture developmental variability.

#### Image segmentation, Feature Selection

For the ABCD study, we accessed raw T1-weighted MRI scans and preprocessed them using FreeSurfer v7.4.1. For UK Biobank, we accessed image derived phenotypes directly as provided by the study resource.

For both data sets we created the same list of 201 brain features, comprising brain volumes of 66 cortical and 135 subcortical regions, using the Desikan-Killiany Atlas. Supplemental Table 1 depicts a list of all features.

#### Harmonization

Since both, the ABCD study and the UK Biobank are multi-site studies, we harmonized imaging data for site confounds using Combat. Specifically, for the longitudinal analysis, we harmonized the data using *LongCombat* package (version 0.0.0.9; ^23^, while for cross-sectional analyses, we used *NeuroCombat* (version 1.0.14; ^24^.

#### Statistical analysis

The statistical analyses were performed using R version 4.4.1. We first used the cross-sectional approach to identify potential overlap between the two hormonal transition phases. Subsequently, we examined whether patterns observed in the cross-sectional analysis would also emerge in the longitudinal analysis by comparing the cross-sectional with the longitudinal results. For all analyses, false discovery rate (FDR) correction was used to correct p-values for multiple comparison ^25^ across all brain volumes.

##### Cross-sectional analysis

For the cross-sectional analyses we used the linear model function *lm* as available in R. For puberty, we included menarche status, age in months, Euler number and site as covariates. The final model was written as:

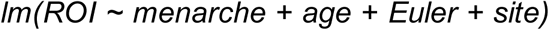

For menopause, menopausal status, age in months, head motion and sites were chosen as covariates, resulting in the following model:

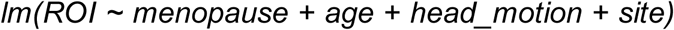

##### Longitudinal analysis

Linear mixed effect (LME) models were fit using the *lme* function of the nlme package (version 3.1.167) in R.

The puberty analysis model included menarche status as the main variable of interest, while age in months, Euler number (image quality proxy), session number (one of three possible time points), and scan sites were included as covariates. Sites with less than 10 observations were dropped. This resulted in the following model:

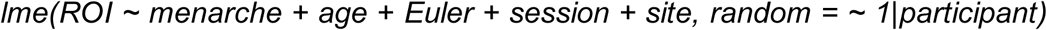

The menopause analysis model contained menopausal status with age in months, head motion and sites as covariates, resulting in the following model:

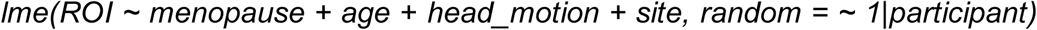

Session was not included as a covariate in the menopause model given the perfect collinearity of session and menopause status.

To address variability from repeated measures, we included participant as a random effect for both models.

Subsequently we examined the descriptive statistics of the cross-sectional results and applied a type II ANOVA from the car package (version 3.1.3) to the longitudinal output, to test the main effects of the menarche and menopause.

## Results

### Cross-sectional Analysis

#### Menarche

For the cross-sectional ABCD dataset, we analysed one linear model per brain region, each accounting for age, Euler number, session and site. We found significant associations with menarche onset for 83 out of 201 brain volumes after correcting for multiple comparisons. Forty-seven of these are cortical and 36 subcortical brain volumes (Supplementary Table 1).

#### Menopause

Linear models on the cross-sectional UK Biobank dataset, resulted in 17 brain volumes significantly affected by the transition into menopause, one cortical and 16 subcortical volumes (Supplementary Table 1).

#### Overlapping regions between cross-sectional approach for Menarche and Menopause

When comparing the 83 brain regions significantly affected by menarche onset with the 17 regions significantly affected by menopause onset in the cross-sectional analysis, seven brain regions overlapped (Figure 1A).

**Figure 1.**
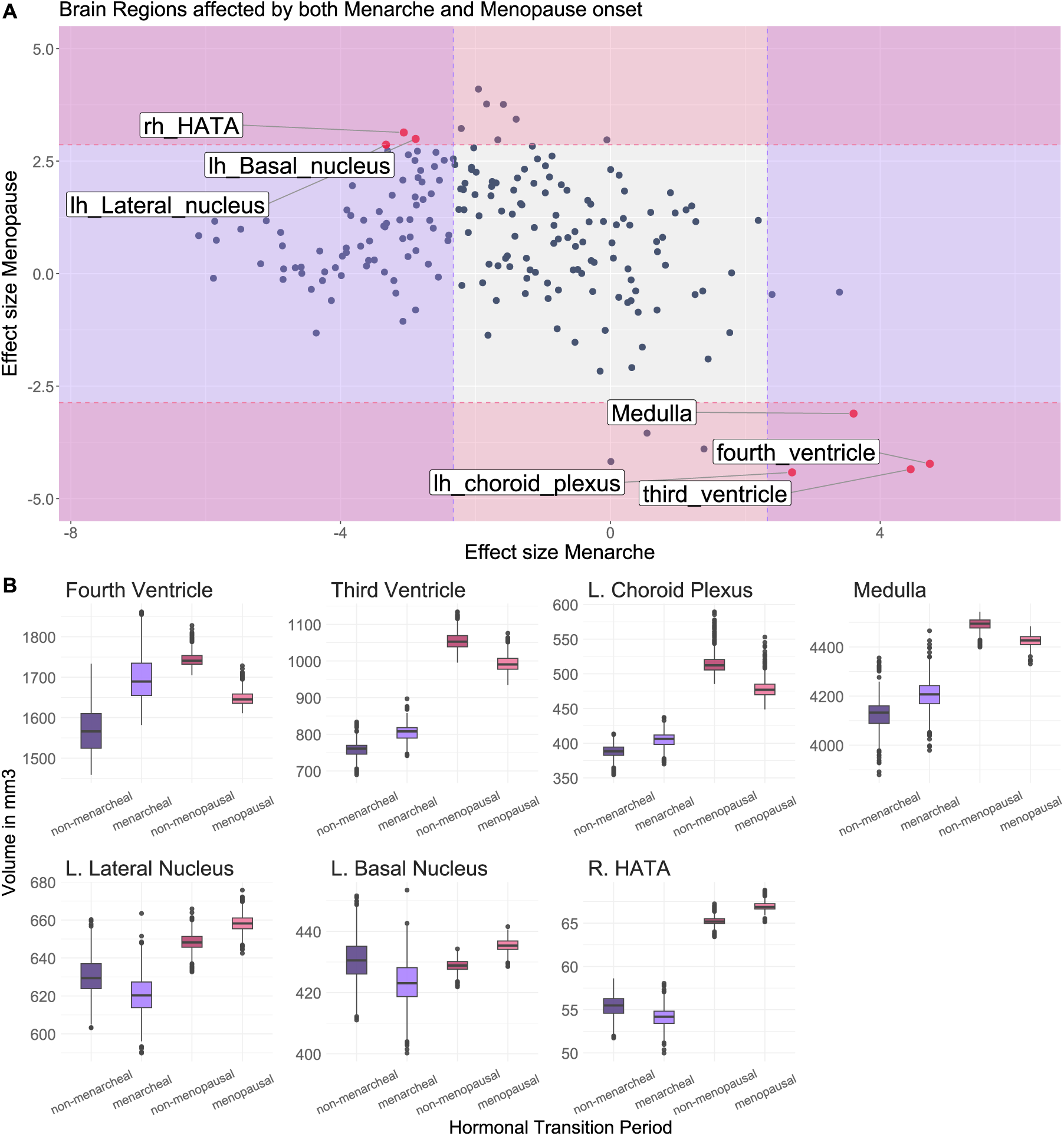
Brain regions affected by both, menarche and menopause. A The scatterplot shows the effect sizes (t-values in mm^3^) of both phases. In red and labelled, are the brain regions that are significantly affected by both transitional phases (passing FDR correction). The dashed lines show the FDR-thresholds. B The boxplots show the volume changes, of the predicted volumes when accounting for effect, age, image quality and site using the predict() function. The purple tones represent the changes during menarche and the red tones during the transition into menopause. L=Left, R=Right.

Furthermore, across all brain regions we observed a weak but statistically significant negative correlation between cross-sectional menarche effects and cross-sectional menopause effect association statistics (Pearson r = −0.14, p = .049, df = 199).

Looking into the direction of the volume changes we observed opposite volume differences between menarche and menopause. Specifically, the fourth ventricle, third ventricle, left choroid plexus and medulla show bigger volumes when starting the reproductive years – menarcheal and not menopausal – compared to the non-reproductive phase. The amygdala regions - left lateral nucleus, left basal nucleus, and right hippocampal amygdala transition area (HATA) show bigger volumes outside the reproductive phase – non-menarcheal and menopausal (Figure 1B).

In an additional analysis, we excluded participants reporting hormonal replacement therapy (HRT). When comparing the 7 regions that were significantly overlapping between menarche and menopause in the main analysis (cross-sectional), 3 volumes remained significantly affected by both phases after controlling for HRT - the fourth ventricle, third ventricle and the left choroid plexus (supplementary figure S3).

### Longitudinal analysis

For each of the two longitudinal datasets (menarche and menopause), we fit linear mixed effect models to each brain volume, accounting for age, Euler number, session and site.

#### Menarche

We found 56 brain volumes significantly affected by menarche onset longitudinally (p < .05 FDR corrected). Of these 56 brain volumes, 35 are cortical and 21 are subcortical regions.

#### Menopause

None of the longitudinal menopause models passed significance threshold for FDR correction. Nine of 201 brain regions were below nominal p level (p<.05; unadjusted). The absence of significance likely reflects the limited statistical power from the small sample size (N=106 per group). Thus, further longitudinal analysis of the effect of menopause onset was not pursued in this study.

#### Convergence between longitudinal and cross-sectional analysis

For the ABCD sample (menarche), 82% of the regions identified as significantly affected longitudinally were also significant cross-sectionally, while 55% of those identified cross-sectionally were identified longitudinally. The overlapping set between both analyses comprised 46 brain volumes – 28 cortical and 18 subcortical (Figure 2A). Furthermore, across all brain regions we observed a strong positive correlation (Pearson r = 0.67, p < 2.2e-16, df = 199) between longitudinal and cross-sectional effect sizes.

**Figure 2.**
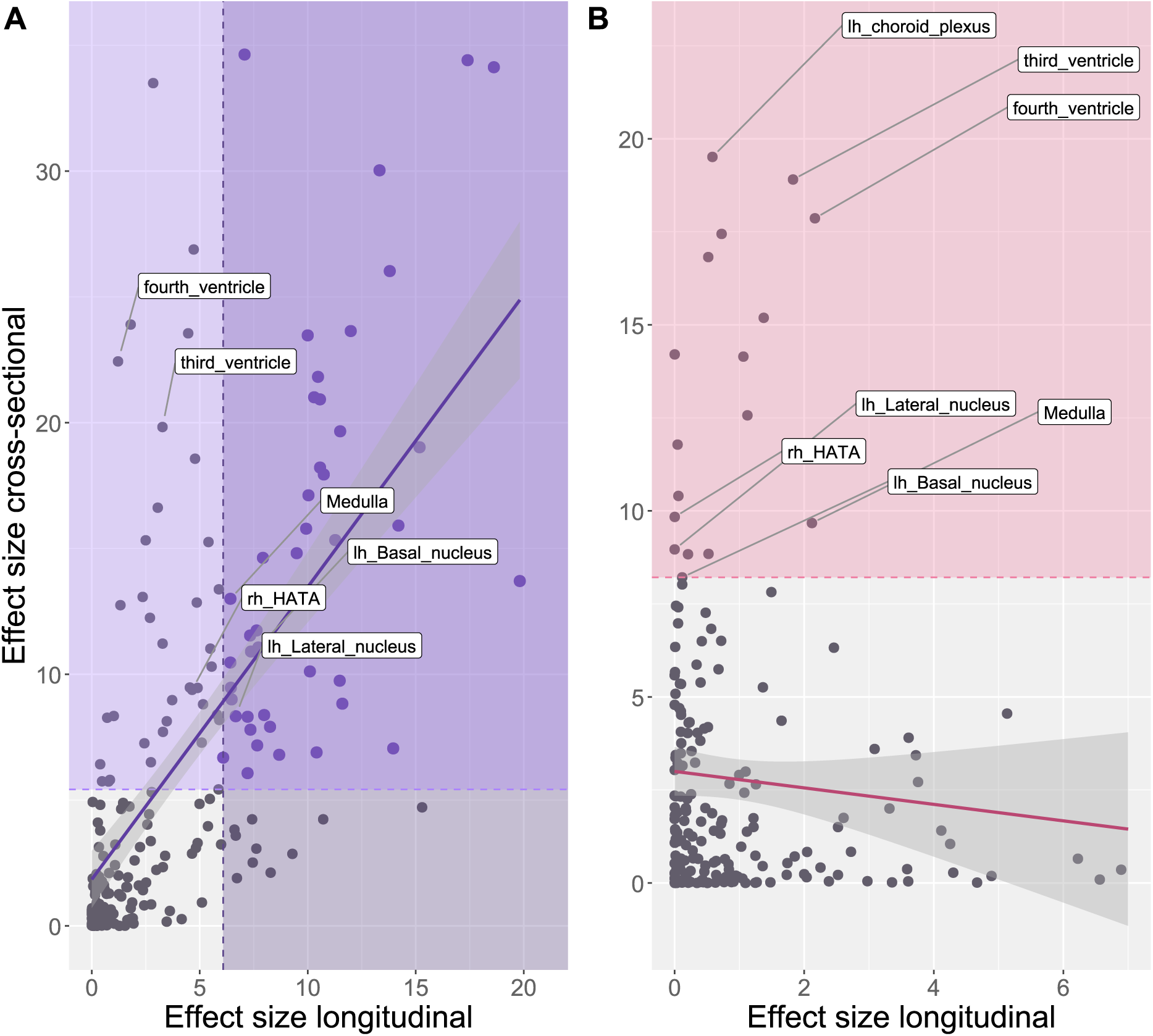
Convergence between the longitudinal and cross-sectional approaches. A Brain structures affected by Menarche. B Brain structures affected by Menopause. For the cross-sectional effect size (in mm^3^), the squared t values and for the longitudinal effect size, the Chisq-values are shown. Labelled are the regions that are significantly affected by both menarche and menopause. The dashed lines show the FDR thresholds.

For the menopause data, a comparison between the cross-sectional and longitudinal results was not possible due to the lack of brain volumes surviving the FDR correction in the longitudinal analysis. A Pearson correlation between the two approaches suggests no significant relationship between the association statistics for the given limited sample size (r = −0.10, p = 0.17, df = 199).

## Discussion

The aim of this study was to investigate and compare brain structural changes during two distinct phases of the female reproductive lifespan: menarche, as a proxy for puberty onset and thus the start of the female reproductive lifespan, and menopause transition, the end of the female reproductive lifespan. Both phases are characterized by profound shifts in levels of sex steroid hormones, especially oestradiol and progesterone, but in opposite directions. Whilst for menarche oestradiol and progesterone levels rise, the transition into menopause shows a notable drop in their levels. We hypothesised that brain regions exist whose volumes are affected by both, puberty and menopause, yet with opposing directions of the effect, in line with what is observed for oestradiol and progesterone levels.

### Overlap between menarche and menopause

Out of the 201 brain volumes that we investigated, we found seven that were significantly affected by both, menarche as well as menopause transition in our cross-sectional analysis. Four of these regions – the fourth ventricle, third ventricle, left choroid plexus and medulla – showed bigger volumes during the reproductive phase than before or after (pre-menarche, post-menopausal). Such an inverted U-shaped trajectory across the lifespan maps onto the above-described pattern of higher oestradiol and progesterone levels during the reproductive years compared to outside of them. Specifically, we observed larger volumes in females reporting the start of their menstrual bleeding compared to females not yet having experienced their menarche. Similarly, females not yet transitioned into menopause and therefore still being in their reproductive phase show bigger volumes than females in menopause. The remaining three regions – the left lateral nucleus, left basal nucleus, and right hippocampal amygdala transition area (HATA) – exhibited a U-shaped trajectory, with bigger volumes outside the reproductive phase when comparing the beginning and end of the reproductive lifespan.

### Menarche

Apart from the analysis of overlap between the two reproductive periods, our analysis also allowed us to shed further light into the individual reproductive periods. Of the 83 brain volumes that were significantly affected by menarche, 92 % presented a negative sign, pointing towards a predominantly negative effect direction, i.e. decrease in volume, in line with previous studies that indicate a general consensus on the global reduction of gray matter volumes during puberty (for a review look at Rehbein et al., 2021) ^6^. We found two out of the seven overlapping regions to be brain volumes of the amygdala – the left lateral nucleus, left basal nucleus and one hippocampal region, the right HATA (hippocampal amygdala transition area) – that show smaller volumes in females who already reached the developmental milestone of menarche compared to females pre-menarche. This pattern could reflect the process of synaptic pruning which is described as a driver for the overall gray matter volume reduction during puberty ^26^. Given the hormonal fluctuations described during the female reproductive lifespan, the observed alterations may be attributed to endocrine variability. However, the role of sex hormones in driving the gray matter volume reduction during puberty is not fully understood yet. Positive and negative associations have been reported between oestradiol levels and different brain volumes such as increased parahippocampal volume ^27^ and decreased volume of the anterior cingulate cortex and prefrontal gray matter density ^7,28^. The administration of oestradiol to adult females has been shown to impact the plasticity of hippocampal subfields ^29^. These heterogeneous reports lead to the assumption that sex hormones, specifically oestradiol, act region-specific. As part of the limbic system, the amygdala is known to have a high density of oestradiol receptors ^30^, is sensitive to hormonal changes and plays a central part in social and emotional processing ^31,32^. Changes in the neuroanatomy of the amygdala are hypothesized to be involved with the heightened vulnerability to mood and anxiety disorders, which emerge particularly in females during and after puberty ^33^. Thus, the observed reduction of amygdala-related brain volumes might correspond to increased exposure to oestradiol levels following menarche onset, leading to region-specific reorganization.

Among the 83 brain regions affected by menarche in the cross-sectional approach, 8% showed a positive effect direction, i.e. increase in volume. This included four of the seven overlapping regions, which are related to the ventricular system – the fourth ventricle, third ventricle, left choroid plexus and medulla – which display bigger volumes in menarcheal females compared to females who reported not having had menarche. The ventricular system is involved in the production and secretion of the cerebral spinal fluid (CSF). The choroid plexus is known for its role in the CSF production ^34^, expresses hormone receptors that can modulate the composition of CSF ^35^ and thereby participates in neuroendocrine signalling ^36^. The observed increased ventricular volumes, similar to the ones described during gestation ^37^, a period marked by increasing steroid hormone levels, might reflect hormonally influenced alterations in CSF regulation which could accompany global tissue reorganization during puberty in females. Together, these findings underscore that puberty is characterized by coordinated yet regionally distinct structural changes in the brain.

Regarding the study design, we found robust convergence of cross-sectional and longitudinal findings, supporting the consistency of our menarche findings. In the cross-sectional analysis, 27% more significant effects were identified compared to the longitudinal approach (for a detailed study design discussion see the Limitations section). Ten regions - three subcortical and seven cortical regions – were significant in the longitudinal but not the cross-sectional menarche analysis, including areas within the prefrontal and cingulate cortex and the accumbens area These regions involved in decision-making and reward processed have been shown to be associated with pubertal development in humans ^38^ as well as animal models ^39^.The discrepancy between the study design approaches may reflect greater inter-individual variability in these regions that is not fully captured by a single time point in the cross-sectional design. Nonetheless, the overall pattern of the cross-sectional results closely aligns with longitudinal ones.

### Menopause

The effect of menopause transition on the 17 significantly affected brain volumes varied in terms of their directions. Fifty-nine % of the brain volumes showed a positive effect, i.e. volume increase following menopausal transition. Regions of the limbic system, including the HATA yield bigger volumes in post-menopausal females compared to pre-menopausal ones, which may initially seem counterintuitive. The menopausal transition has been linked with difficulties in memory and increased risk for Alzheimer’s disease potentially induced by oestrogen deficiencies in older females ^40,41^. Therefore, a large proportion of studies investigating neuroplasticity during menopausal transition primarily examined the neuroprotective effect of menopause hormonal therapy (MHT) with a particular focus on hippocampal regions ^42–44^. However, studies investigating specifically the effect of menopausal transition itself yielded inconsistent findings. Some are reporting a decline of hippocampal volume in post-menopausal females compared to pre-menopausal females ^45,46^ and others are reporting no significant influence on hippocampal volumes based on menopause status ^47^. Differences in sample size (N = 33 in Sullivan et al. ^47^ and N= 108 in Goto et al. ^46^) may partly explain the discrepant findings as well as the use of MHT, as Sullivan et al. included MHT users, while Goto et al. did not, suggesting that hormonal therapy may mask menopause-related effects on hippocampal structures. However, other studies reported no volumetric differences comparing MHT users and non-users ^48^, emphasising the complexity of these effects. Furthermore, detangling the effects of chronological aging from the effects of menopause remains challenging. To address this issue, Mosconi et al. compared females in different menopausal phases to age-matched men, concluding that observed brain changes reflect menopausal aging rather than chronological aging ^49^. Additionally, in a two-year follow-up session, they found partial reversal of brain regions relevant to cognition comparing in the post-menopausal females. Our results align with these findings, underscoring the sensitive interplay of hormonal status and aging.

Forty-one percent of the 17 significantly affected brain volumes from the cross-sectional menopause analysis showed a negative effect, i.e. a volume reduction. Amongst these, we observed four regions of the ventricular system to be affected by menarche and menopause. They presented bigger volumes in pre-menopausal females compared to post-menopausal. Brain aging is typically accompanied by global volume reduction and corresponding linear CSF volume increase. While pathological ventricular enlargement can be used as a biomarker for Alzheimer’s disease ^50^, it also occurs in normal chronological aging, where increase in ventricular volume tends to be more amplified after the age of 60 years ^51^. In our sample, mean age of post-menopausal females was 55 years, suggesting this age-related increase might not have occurred yet. The observed volume reduction mimics the pattern of steroid hormones, which decrease with the transition into menopause. Knowing that the choroid plexus entails receptors for steroid hormones, there might be a link, however, the underlying mechanism remains unclear. Our findings indicate that fluctuations in CSF production might contribute to brain volume dynamics during transitional phases, indicating that CSF-mediated mechanisms might play a more important role in neuroplasticity during endocrine changes.

Investigating the convergence between cross-sectional and longitudinal design was not possible, as no effect survived correction for multiple comparison presumably reflecting the limited longitudinal sample size.

### Limitations

Another aim of our study was to compare different study design approaches. Menarche and menopause are highly age-dependent biological processes. Consequently, approaching these cross-sectionally by comparing age-matched participants that have already gone through the process with those who have not, one could argue this may lead to selection bias such as for example selecting people with atypical timing for one of the groups. Selection bias can have large implications for interpretations. In our study, we included only participants within a typical age range for puberty or menopause onset. This could result in controls who have not experienced menarche or menopause within that plausible age range, thus being atypical by design. Consequently, unbiased selection of controls is challenging unless longitudinal data is available to track the timing, enabling more comparable grouping. Additionally, there might be baseline differences in participants of the cross-sectional analysis that we cannot account for. To reduce this bias, we sampled from a very limited age range, incorporating this constraint into the matching process. Moreover, we applied a longitudinal approach in addition to our cross-sectional design. For the menarche groups, we observed a strong convergence between results of the cross-sectional and longitudinal analysis, supporting the robustness of our results. Thus, a longitudinal design provides valuable means to minimize selection bias and might generally be better in capturing within-subject changes and individual trajectories while the cross-sectional approach serves as a valuable tool for screening for effects. However, longitudinal studies are resource-intensive and therefore harder to implement, which was notable for our menopause data, where we were not able to test the convergence, because of the lack of longitudinal sample

We chose menarche as the final physical event of female pubertal development for our analysis to be able to compare individuals before and after this biologically critical event. Given that the scans in the ABCD are approximately two years apart from each other, and the self-reported date of menarche partially varies a lot between the children’s and caregiver’s report, the precise timing of menarche and start of normal cycle is not known. Consequently, the difference between menarche and the closest scan could vary across participants. Nevertheless, we assumed, that this variability is evenly distributed across the number of participants included in the analysis. Similarly, menopause is diagnosed retrospectively – cessation of menstruation for 12 consecutive months ^52^– which makes the self-report and exact timing of start of menopausal transition complex. These parallel limitations highlight the need for more longitudinal studies focused on pubertal and menopausal timing and the incorporation of more female-specific questions in aging studies to enhance accuracy and reliability.

We did not account for the use of hormonal contraceptives in the postmenarcheal and premenopausal females, which would change the hormonal fluctuations by suppressing ovulation and thereby have an impact on the neuroplasticity ^53,54^ and therefore the interpretation of our findings. However, when controlling for another form of hormonal intervention – menopause hormone therapy, please see supplemental analysis – several regions remained significantly affected by the transition, pointing towards not all significant results are attributable to hormonal interventions.

Finally, the absence of hormonal data in this study poses a limitation to the interpretation of our findings as a direct comparison of brain structural changes and hormone level changes is not possible. Menarche and menopause are both processes that are highly driven by hormonal changes. We here mostly discuss the influence of oestradiol, but other hormones like testosterone also fluctuate across the lifespan and have been shown to influence female neuroplasticity ^55^. The observed brain structural changes might partly be explained by neuroendocrine changes but other factors such as water homeostasis and socio-economics cannot be neglected. To further investigate the influence of female specific hormones, introducing a male control group could offer some valuable comparative insights. However, this would also impose some challenges as menarche and menopause are both female specific processes and pubertal onset offers later in males, which would complicate the age-matched comparison.

## Conclusion

This study demonstrates that the female brain undergoes substantial neural plasticity both at the onset and the ending of the reproductive lifespan. The study revealed several brain regions, including limbic and ventricular regions, being affected by both menarche and menopause. As hypothesized, overlapping regions showed opposite effect directions between menarche and puberty. The significant overlap was smaller than expected, underscoring that factors like socioeconomic status may play crucial roles alongside hormonal fluctuations. Overall, these findings highlight the importance of examining (endocrine-driven) neuroplasticity not only during individual transitions but also across the whole female lifespan in order to better understand the overarching mechanisms. Due to the complexity of these phases, mainly driven by hormonal fluctuations, further research would benefit from the incorporation of hormonal assessments and extension of the investigation to other transitional phases like pregnancy, a critical period of rapid hormonal transition and brain remodelling.

## Acknowledgement & Funding

T.K. received funding from the European Research Council (ERC CoG, #101086793, HealthyMom), and the Research Council of Norway (#323961). G.M.B., A.S., B.D. and T.K. were supported by the German Research Foundation (IRTG 2804). Data analysis was supported by the BMBF-funded de.NBI Cloud within the German Network for Bioinformatics Infrastructure (de.NBI) (031A537B, 031A533A, 031A538A, 031A533B, 031A535A, 031A537C, 031A534A, 031A532B).

## Data and Code availability

Both data sets were accessed under data use agreements with the data providers.

The ABCD Study® (https://abcdstudy.org), held in the NIMH Data Archive (NDA), is a multisite, longitudinal study designed to recruit more than 10,000 children age 9-10 and follow them over 10 years into early adulthood. The ABCD Study® is supported by the National Institutes of Health and additional federal partners under award numbers U01DA041048, U01DA050989, U01DA051016, U01DA041022, U01DA051018, U01DA051037, U01DA050987, U01DA041174, U01DA041106, U01DA041117, U01DA041028, U01DA041134, U01DA050988, U01DA051039, U01DA041156, U01DA041025, U01DA041120, U01DA051038, U01DA041148, U01DA041093, U01DA041089, U24DA041123, U24DA041147. A full list of supporters is available at https://abcdstudy.org/federal-partners.html. A listing of participating sites and a complete listing of the study investigators can be found at https://abcdstudy.org/consortium_members/. ABCD consortium investigators designed and implemented the study and/or provided data but did not necessarily participate in the analysis or writing of this report. This manuscript reflects the views of the authors and may not reflect the opinions or views of the NIH or ABCD consortium investigators. The ABCD data repository grows and changes over time. The ABCD data used in this report came from https://doi.org/10.15154/zk5y-pc91.

UK Biobank data was accessed under the application number #121699.

Analysis code will be made publicly available at GitHub upon publication.

## Authors Contribution

M.F. Conceptualization; Formal Analysis; Methodology; Software; Visualization; Writing – original draft (lead); Writing – review and editing

G.M.B Methodology; Software; writing – review and editing

A.S. Supervision, Writing – review and editing

B.D. Supervision, Writing – review and editing

T.K. Conceptualization, Funding Acquisition, Methodology, Supervision, Writing – original draft (supporting); Writing – review and editing

## SUPPLEMENT

**S1.**
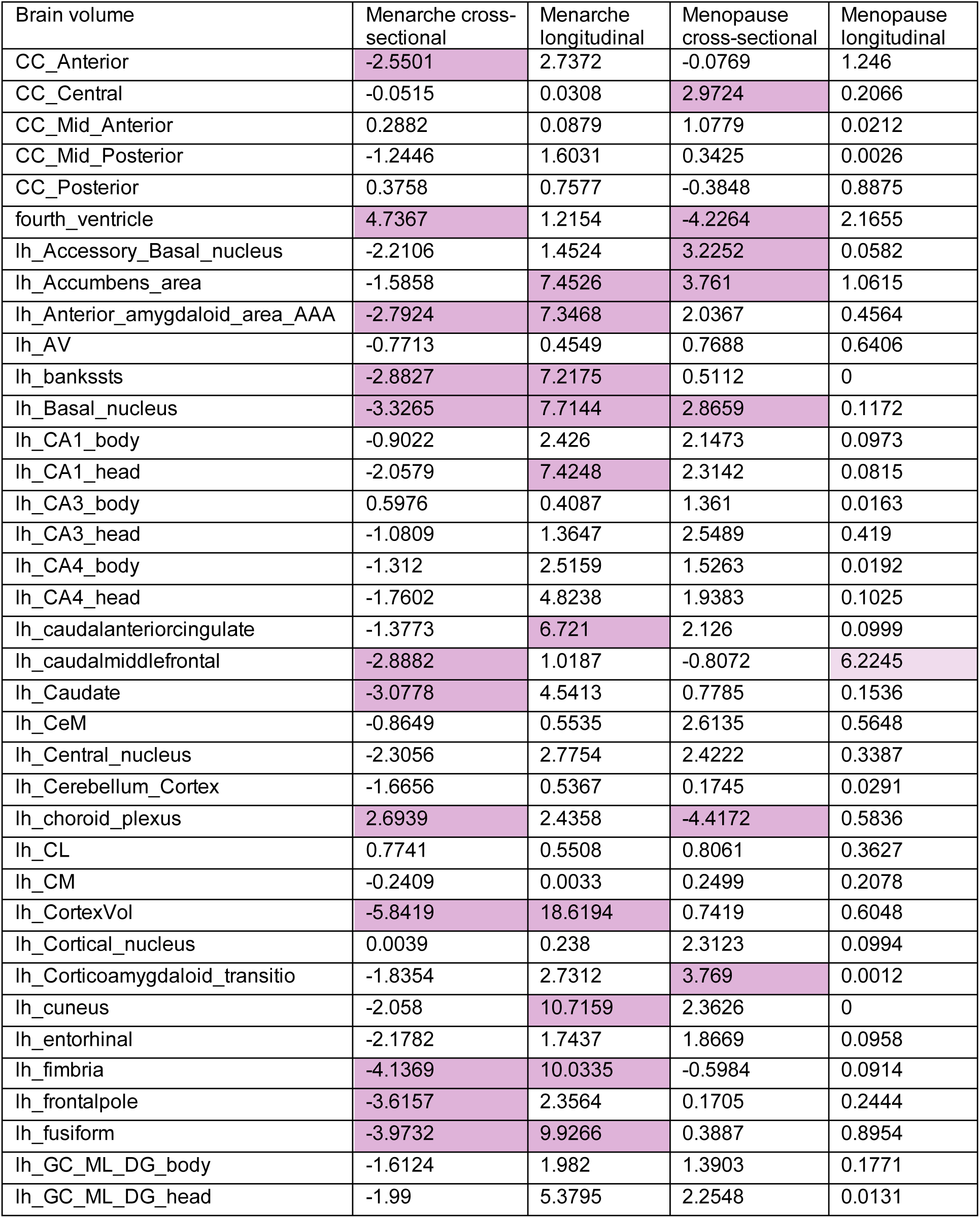
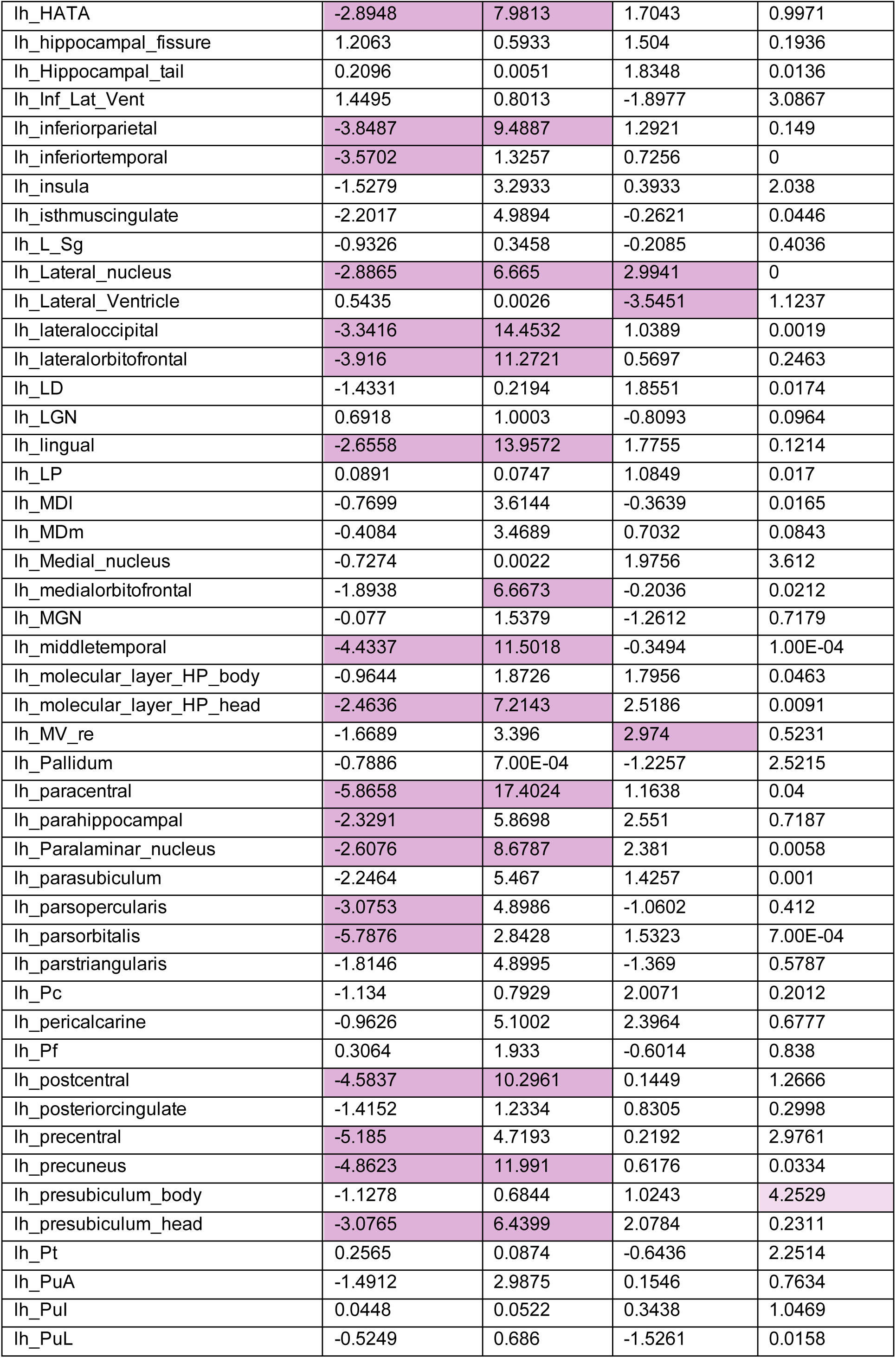
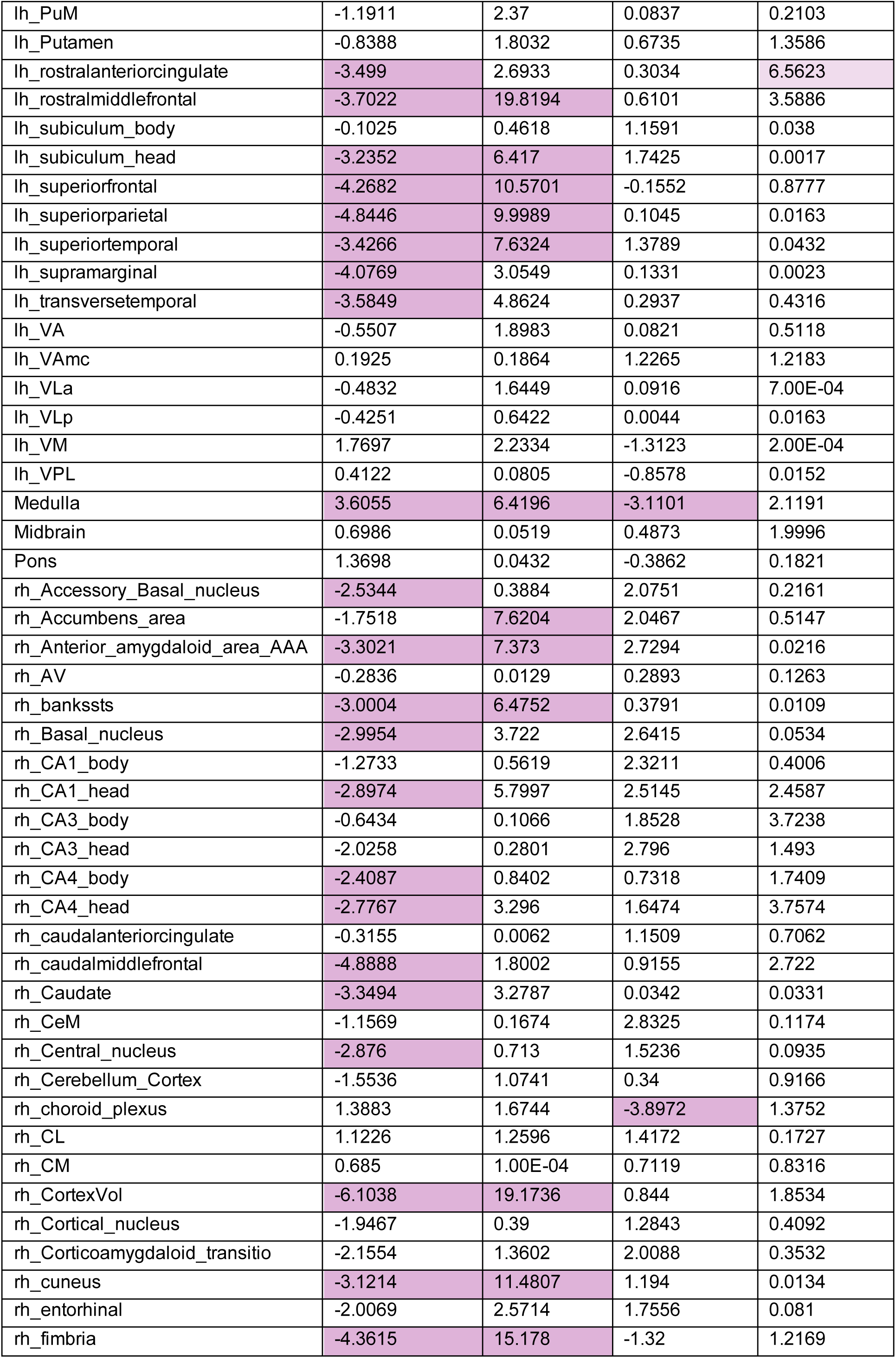
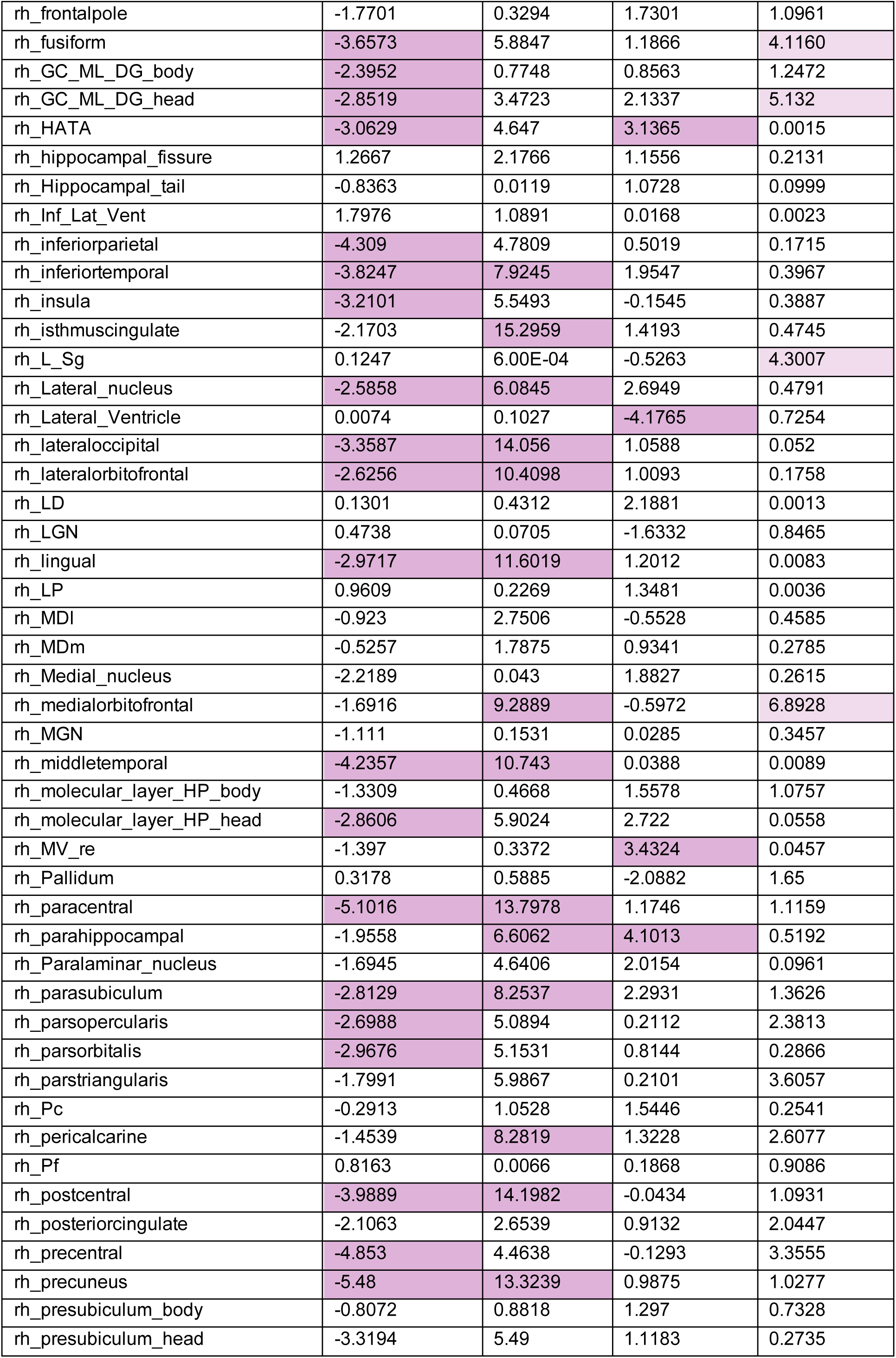
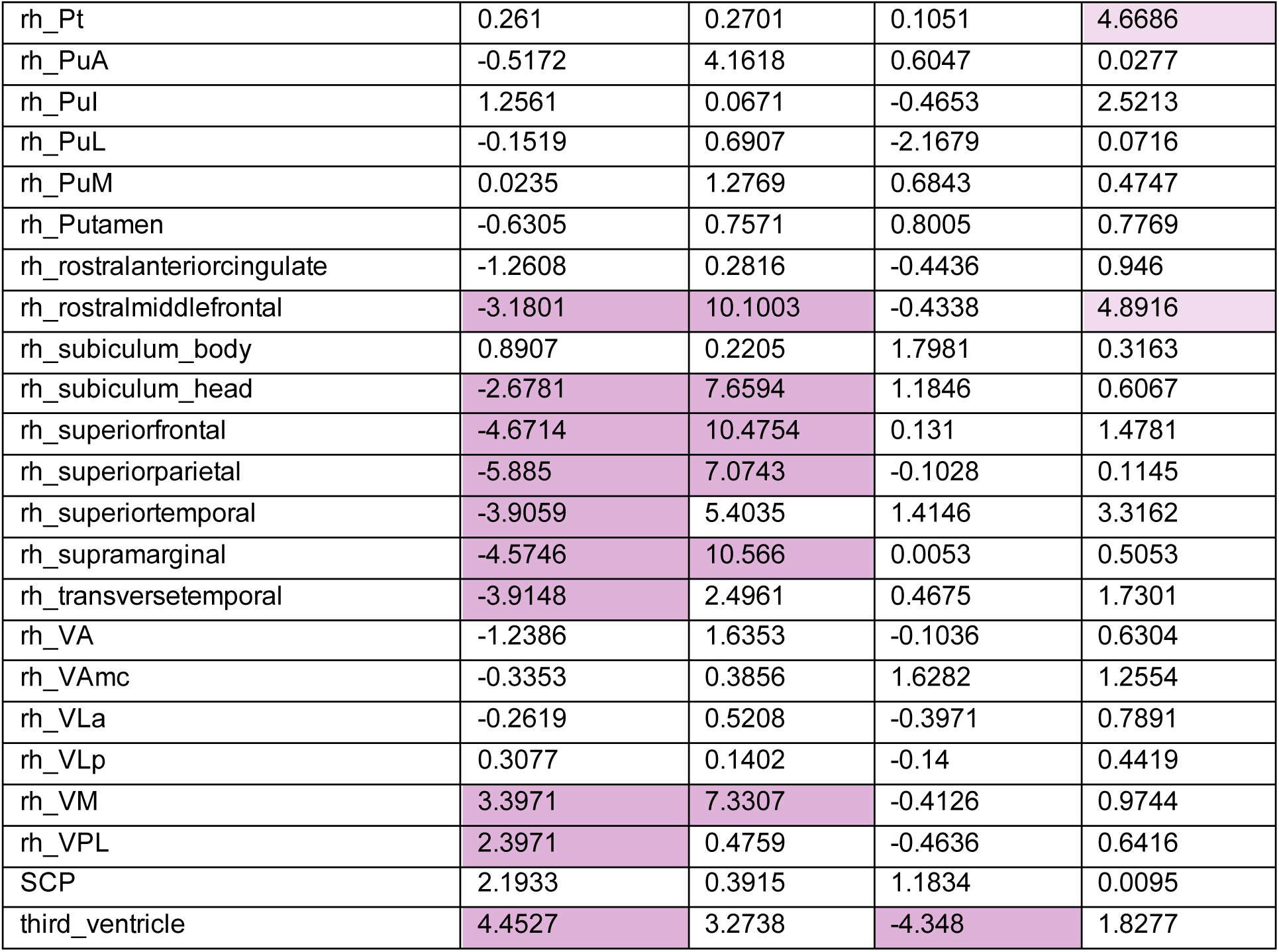
Brain volumes used for the analysis and the according effect sizes for each analysis. For the cross-sectional analysis, t-values and for the longitudinal analyses, Chisq-values are reported. The significant FDR thresholds for the effect sizes are as followed: for longitudinal menarche = 6.0845, cross-sectional menarche = +/− 2.329, cross-sectional menopause = +/− 2.8659. Significant values are coloured in purple. For the longitudinal menopause analysis no brain volume survived the correction for multiple comparison, the lighter purple shows the effect sized that exceeded the nominal significance threshold of 4.116.

### Comparison of including and excluding participants with Menopause Hormone Therapy

To rule out potential confounds due to menopause hormone therapy, we run supplementary analysis on the menopause dataset, excluding participants reporting the use of menopause hormone therapy. For the longitudinal analysis this resulted in 79 participants pre-menopause (mean age = 51.3 years, sd = 2.3) for which we have follow-up data in the menopausal group (mean age = 55.2 years, sd = 2.3).

For the cross-sectional analysis, we excluded participants reporting menopause hormone therapy based on the main cross-sectional sample. This resulted in a sample size of 1146 participants – 573 non-menopausal (mean age = 52.7 years, sd =2.2 years), 573 menopausal (mean age = 52.8 years, sd = 2.2 years).

We ran a Pearson correlation between the datasets containing MHT participants and the ones, excluding MHT participants. The analysis resulted in a strong positive correlation of r = 0.70 (t = 13.756, df = 199, p-value < 2.2e-16) for the longitudinal dataset as well as for the cross-sectional dataset with r = 0.77 (t = 17, df = 199, p < 2.2e-16; Figure S2).

Additionally, we compared the cross-sectional menopause dataset, excluding participants reporting MHT intake and the cross-sectional menarche dataset. Three brain volumes, the fourth ventricle, third ventricle and the left choroid plexus remained significantly affected by both transitional periods. A Pearson correlation ran between the two datasets resulted in a weak, non-significant correlation between the two datasets, with r = −0.11 (t = −1.6091, df = 199, p-value = 0.1092; Figure S3).

**Figure S2.**
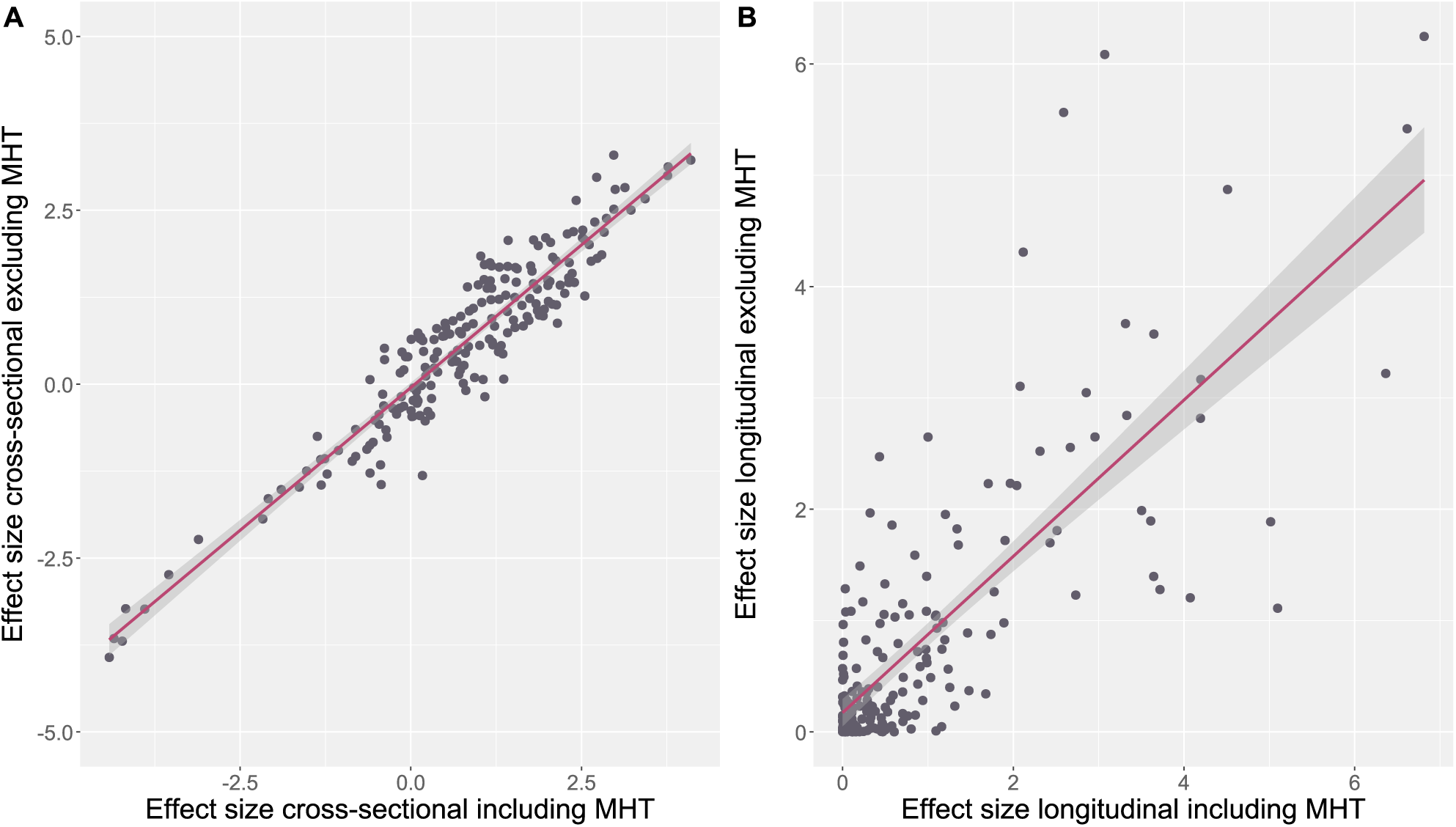
Scatterplot comparing the effect sizes of the Menopause onset including and excluding participants reporting MHT intake. A shows the cross-sectional and B the longitudinal approach.

**Figure S3.**
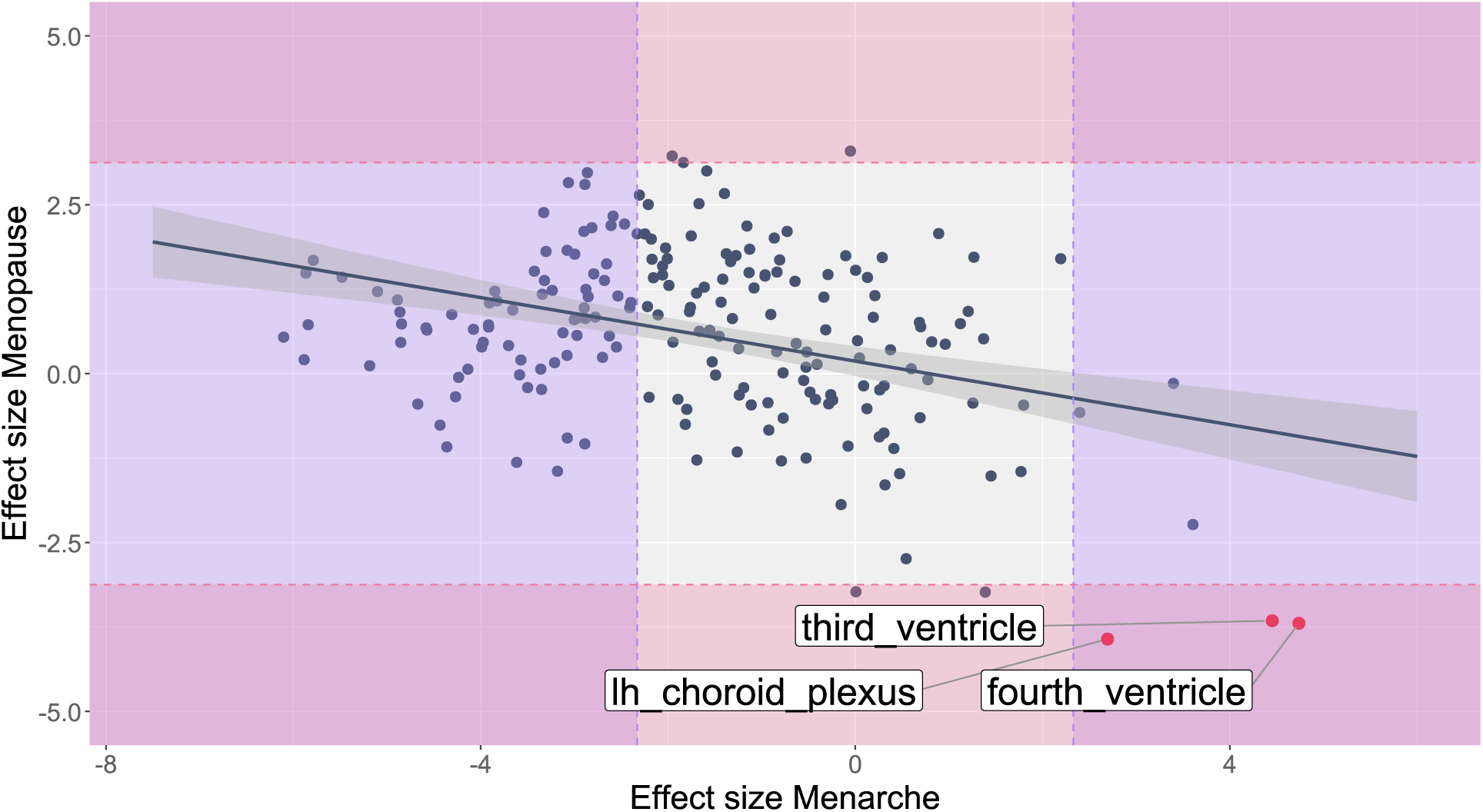
Brain regions effected by both, menarche and menopause (excluding participant reporting MHT intake). The scatterplot shows the effect sizes (in mm^3^) of both phases. In red and labelled, are the brain regions that are significantly affected by both transitional phases. The dashed line shows the FRD-threshold.

